# Mechanical Resistance to Micro-Heart Tissue Contractility unveils early Structural and Functional Pathology in iPSC Models of Hypertrophic Cardiomyopathy

**DOI:** 10.1101/2023.10.30.564856

**Authors:** Jingxuan Guo, Huanzhu Jiang, David Schuftan, Jonathan D Moreno, Ghiska Ramahdita, Lavanya Aryan, Druv Bhagavan, Jonathan Silva, Nathaniel Huebsch

**Affiliations:** Department of Mechanical Engineering and Material Science; Department of Biomedical Engineering; NSF Science and Technology Center for Engineering Mechanobiology; Division of Cardiology, Department of Medicine, Washington University in St. Louis, St. Louis, Missouri, USA; Center for Cardiovascular Research, Center for Regenerative Medicine, Center for Investigation of Membrane Excitability Diseases, Washington University in Saint Louis

**Keywords:** Mechanical loading, MYBPC3, bioengineering, calcium, contractility

## Abstract

Hypertrophic cardiomyopathy is the most common cause of sudden death in the young. Because the disease exhibits variable penetrance, there are likely nongenetic factors that contribute to the manifestation of the disease phenotype. Clinically, hypertension is a major cause of morbidity and mortality in patients with HCM, suggesting a potential synergistic role for the sarcomeric mutations associated with HCM and mechanical stress on the heart. We developed an *in vitro* physiological model to investigate how the afterload that the heart muscle works against during contraction acts together with HCM-linked MYBPC3 mutations to trigger a disease phenotype. Micro-heart muscle arrays (μHM) were engineered from iPSC-derived cardiomyocytes bearing MYBPC3 loss-of-function mutations and challenged to contract against mechanical resistance with substrates stiffnesses ranging from the of embryonic hearts (0.4 kPa) up to the stiffness of fibrotic adult hearts (114 kPa). Whereas MYBPC3^+/-^ iPSC-cardiomyocytes showed little signs of disease pathology in standard 2D culture, μHMs that included components of afterload revealed several hallmarks of HCM, including cellular hypertrophy, impaired contractile energetics, and maladaptive calcium handling. Remarkably, we discovered changes in troponin C and T localization in the MYBPC3^+/-^ μHM that were entirely absent in 2D culture. Pharmacologic studies suggested that excessive Ca^2+^ intake through membrane-embedded channels, rather than sarcoplasmic reticulum Ca^2+^ ATPase (SERCA) dysfunction or Ca^2+^ buffering at myofilaments underlie the observed electrophysiological abnormalities. These results illustrate the power of physiologically relevant engineered tissue models to study inherited disease mechanisms with iPSC technology.

## Introduction

Hypertrophic cardiomyopathy is the most common inherited cardiomyopathy and the leading cause of sudden cardiac death in the young. The incomplete penetrance of HCM(1), for example, HCM patients from the same family, and even twins, harboring identical sarcomere genomic variants can develop vastly different phenotypes (2, 3). Thus, non-genetic, environmental factors such as hypertension and associated myocardial stiffening may play a synergistic role in developing an overt HCM phenotype(4–6). Hypertension causes increased afterload and mechanical loading on heart muscle which induces cardiac hypertrophy. How HCM mutations and mechanical stress on the heart might synergize to unmask a pathologic phenotype of HCM remains unknown(7). Advances in tissue engineering and stem cell biology have improved structural and functional maturity of induced pluripotent stem cell (iPSC) derived cardiomyocytes, making these cells valuable tools for studying inherited heart disease(8–11). However, unlike mouse models where transverse aortic constriction has been used to study pressure overload effects on disease pathophysiology(6), there have been few *in vitro* engineered heart tissue-based studies that investigate how HCM mutations alter cardiomyocyte hypertrophic remodeling in response to mechanical loading(4, 5).

HCM is linked to genetic mutations of protein components of the contractile sarcomere apparatus of cardiomyocytes. Mutations in myosin binding protein C (MYBPC3) predominate, the vast majority of which are loss-of-function mutations(12–15). Hallmarks of the HCM phenotype include exuberant myocyte hypertrophy, structural disarray, and the propensity for malignant cardiac arrhythmia. Current medical management including beta-blockers, calcium channel blockers, and the novel myosin inhibitor, mavacamten, focus on treatment of symptoms(7). Interestingly, valsartan, an afterload reducing agent, has been found to inhibit the progression of the HCM phenotype, but only when the drug was given before the HCM phenotype manifested in animal models, suggesting that afterload may play a critical role in disease manifestation(16, 17). Nevertheless, the precise molecular mechanisms through which these drugs can reduce and potentially prevent hypertrophic remodeling and/or abnormal calcium handling is not clear(18, 19).

Clinical and animal studies of HCM suggest that cellular and sarcomeric disarray within cardiomyocytes may be directly related to the sarcomere mutation, with fibrosis and electrophysiological abnormalities being secondary(20, 21). In iPSC models, sarcomere disarray was reported with MYH7 mutations. However, no clear sarcomere disarray was seen in MYBPC3^-/-^ cardiomyocytes(5, 15, 22–24). In addition to structural disorganization of the sarcomere, altered Ca^2+^ intake and increased Ca^2+^ sensitivity is often described in HCM(25–28). For example, analysis of cardiomyocytes from HCM patients’ surgical biopsies has shown reduced expression of sarcoplasmic reticulum Ca^2+^ ATPase (SERCA) and inefficient contractile energetics(29). Studies using iPSC models have identified abnormalities in the Ca^2+^ transient with contractile deficits(5, 27). However, iPSC models also identified sarcomere mutation induced contractile dysfunction, which were independent of Ca^2+^ signaling(15, 22, 23). Thus, it remains unclear whether Ca^2+^ dysfunction in the setting of HCM is triggered by environmental factors.

Here, we investigated how mechanical resistance to contractility, together with a physiologically relevant aligned 3D culture environment, impact early development of HCM phenotypes in MYBPC3^+/-^ and isogenic control iPSC-derived micro-heart muscle arrays (μHM). We hypothesized that increased resistance to contractility in the form of substrate stiffness would mimic the effects of hypertension induced afterload to trigger early pathogenesis of HCM. To provide this mechanical resistance, we formed μHM on poly(dimethylsiloxane) (PDMS) substrates with elasticity (tensile young’s moduli) ranging from 0.4 kPa to 114 kPa to mimic myocardial stiffnesses ranging from the embryonic developmental stage up to a fibrotic adult heart muscle(20, 30, 31)(**Fig. 1**). Within µHMs, MYBPC3^+/-^ cardiomyocytes exhibited cellular hypertrophy, along with changes in sarcomere morphology, contractility, and calcium handling which were absent in 2D monolayers. The changes were exacerbated by increasing the stiffness of the substrate that µHMs contracted against. Pharmacologic studies pointed towards excessive calcium intake through *L*-type channels, rather than SERCA dysfunction or calcium buffering at the myofilaments, as the most likely explanation for aberrant Ca^2+^ transients and contractility. We also identified some micro-scale deficits in troponin localization in MYBPC3^+/-^ tissues in the setting of early stage HCM pathogenesis. These studies suggest that early initiation of the HCM phenotype may involve micro-scale sarcomere reorganization, which depends on afterload sensitivity, emphasizing the value of engineered tissue models in uncovering pathogenic mechanisms in the early stages of HCM.

**Figure 1.**
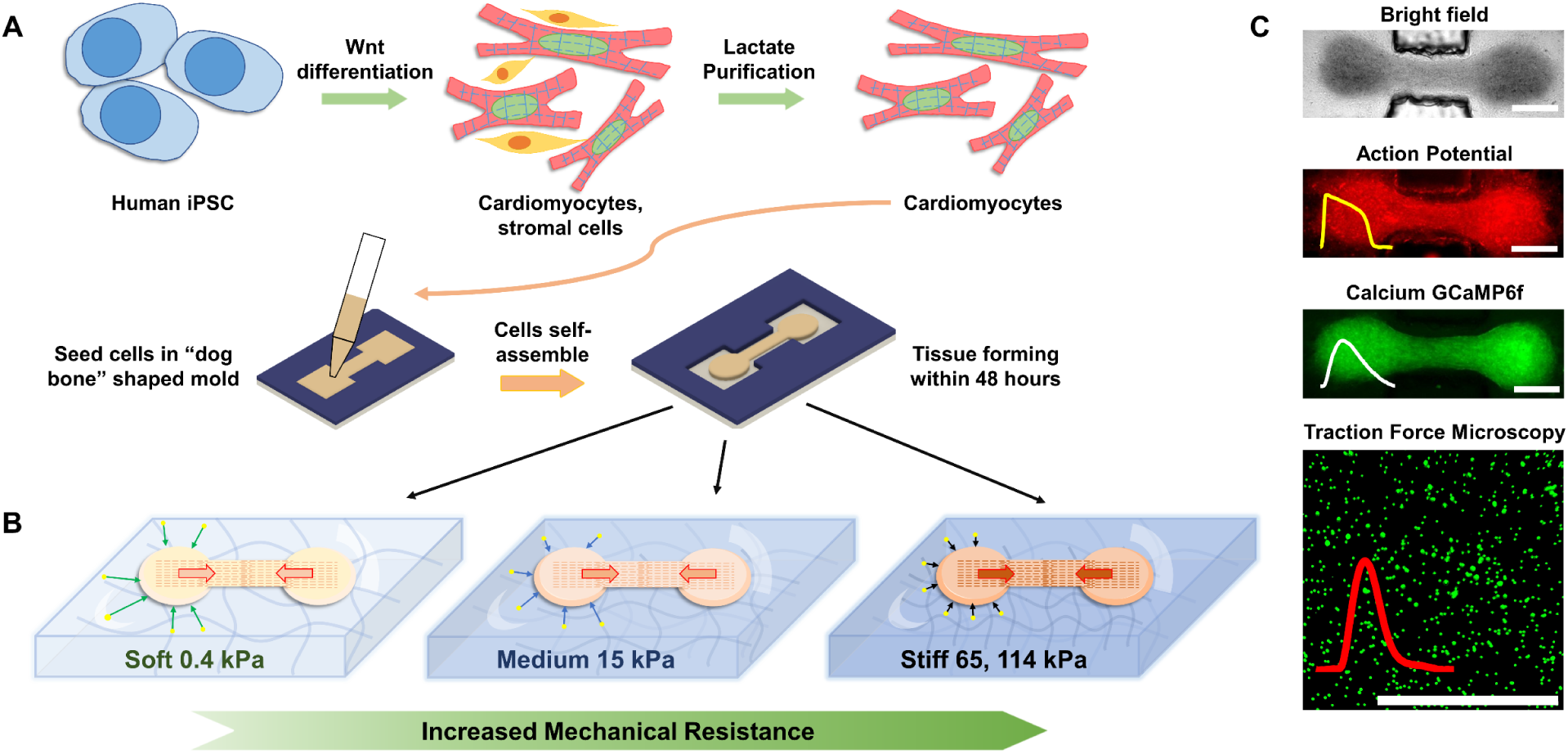
Schematic of mechanical overload enhanced system. (A) Human iPSC differentiation using Wnt signaling pathway, and cardiomyocytes were metabolically purified using lactate. High density of cardiomyocytes mixture was filled into “dog bone” shaped stencil mold to form tissue. (B) Tissue within “dog bone” shaped mold was formed on PDMS substrates with desired mechanical stiffness mimicking different cardiac resistance during contraction. (C) iPSC-µHM action potential, calcium and traction forces were measured through fluorescence high speed imaging.

## Results

### Mechanically optimized µHM substrates reveal HCM genotype-dependent cellular hypertrophy and isoproterenol responses

As HCM does not manifest in utero, our expectation was that substrates mimicking embryonic heart elasticity would not lead to appreciable differences between MYBPC3^+/-^ and isogenic control tissues. Conversely, disease phenotypes would develop in tissues working against substrates with postnatal levels of stiffness. Consistent with this expectation and prior studies(4, 32), there was no evidence for hypertrophy in MYBPC3^+/-^ iPSC-cardiomyocytes in 2D (**SI Figure. 2**). Within μHM, increasing substrate stiffness triggered cellular hypertrophy in both genotypes. However, cryosections of MYBPC3^+/-^ tissues revealed a significant increase in cellular cross-sectional area compared to what was observed in isogenic controls (**Fig. 2 A, B**). This difference was exacerbated on substrates mimicking the stiffness of healthy heart muscle (15 kPa) or hearts at intermediate stages of fibrosis (65 kPa)(31, 33), with MYBPC3^+/-^ cardiomyocytes exhibiting over a 30% increase in projected cell area. In contrast, on very stiff (114 kPa) substrates, cardiomyocytes within control μHM exhibited substantial hypertrophy, blunting genotype-linked differences (**Fig. 2 B**). This result mimics prior observations from both clinical and mouse studies(34, 35), where MYBPC3 deficient mice had similar heart mass compared to wild-type controls at the baseline (no mechanical stress) but exhibited cardiac hypertrophy in response to hypertension caused by transaortic constriction surgery(6).

**Figure 2.**
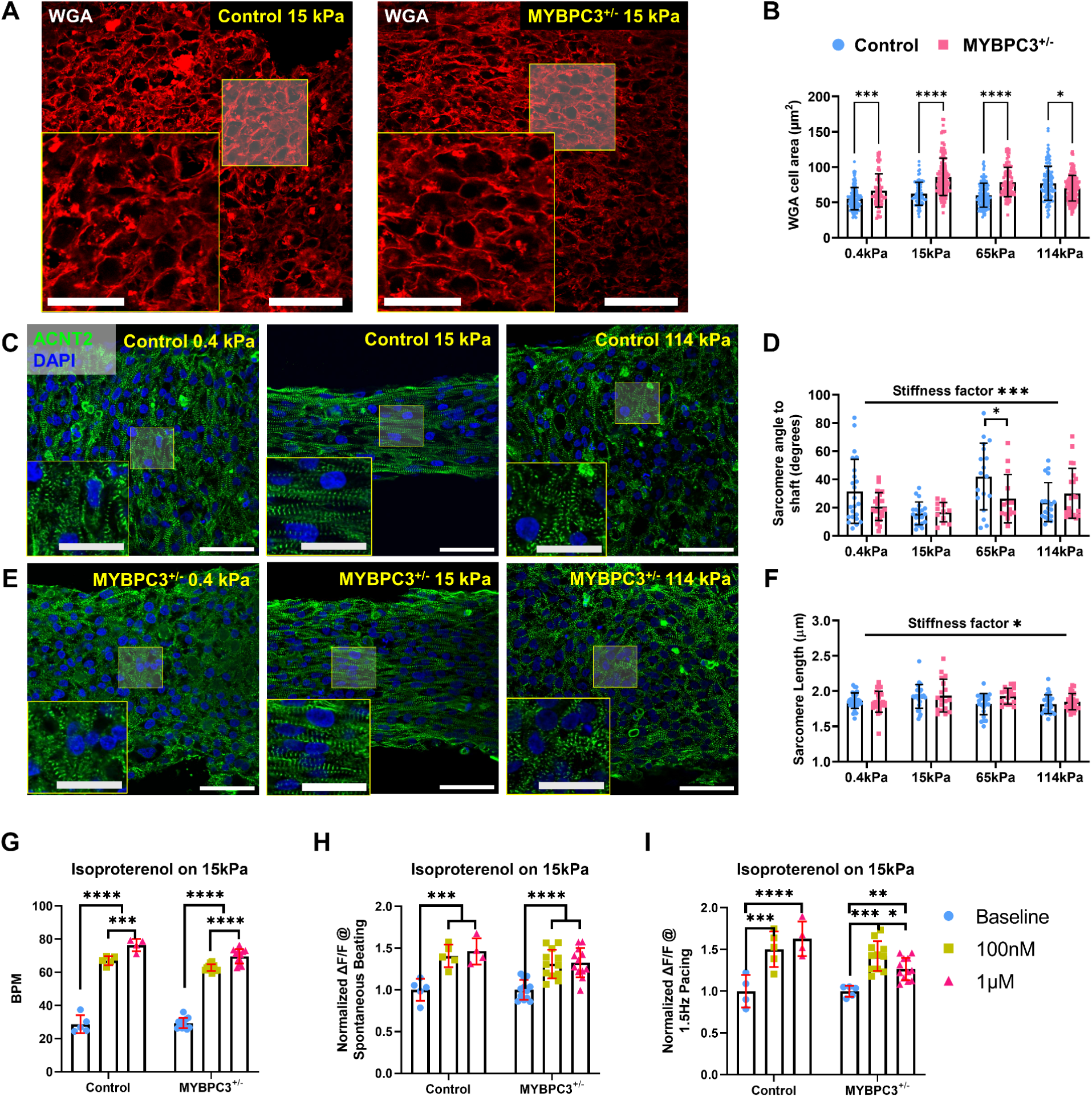
Substrate stiffness trigger iPSC-µHM morphological adaptation and hypertrophy. (A) Representative WGA staining for control and MYBPC3^+/-^ iPSC-µHM at 15 kPa, larger cellular cross-sectional area was seen for MYBPC3^+/-^ tissues. (B) Mechanical stiffnesses trigger cellular hypertrophy. MYBPC3^+/-^ has larger cell area compared to control tissue, especially at 15 and 65 kPa conditions, however, less genotype related hypertrophy was seen at 114 kPa. (C) Representative sarcomere α-actinin and DAPI co-stain indicates both control (top) and MYBPC3^+/-^ (bottom) iPSC-µHM has better sarcomere alignment at 15 kPa condition. Less organized sarcomere was observed at ultrasoft 0.4 kPa and stiff 114 kPa conditions. (D) Average angle between individual sarcomere to tissue shaft. 15 kPa condition has the most organized sarcomere, both genotypes adapt to environmental stiffness similarly. (E) Average sarcomere lengths were similar between genotypes, 15 kPa condition has slightly longer sarcomere compared to other conditions. (F) Spontaneous beat rate changes for control and MYBPC3^+/-^ 15 kPa tissues in response for isoproterenol. (G-H) Spontaneous (G) and 1.5 Hz (H) pacing Ca^2+^ intake for control and MYBPC3^+/-^ 15 kPa tissues in response for isoproterenol. *, ***, **** indicates *p* value less than 0.05, 0.001 and 0.0001. Error bar: *SD*. n represents >3 individual differentiated tissue batches. Scale bar: 50 µm, insets: 25 µm.

Despite the profound cellular hypertrophy in MYBPC3^+/-^ tissues, the overall Z-disk organization, as depicted by staining for sarcomeric α-Actinin (ACTN2) was similar between MYBPC3^+/-^ and control iPSC-cardiomyocytes cultured in 2D on tissue culture plastic and in 3D µHM (**Fig. 2 C-F**; **Fig. 4 B-C**). This observation suggests that hypertrophic remodeling may precede gross cardiomyocyte disorganization and sarcomere disarray in the context of early stage HCM. In both MYBPC3^+/-^ and isogenic control μHM, increasing substrate stiffness from 0.4 kPa to levels reflective of healthy adult myocardium (15 kPa) markedly enhanced cellular alignment and sarcomere orientation in both genotypes, illustrating mechano-induced structural maturation(36). Notably, in μHM formed on stiff 114 kPa substrates, we observed poor sarcomere organization as quantified by a high divergence in sarcomere alignment (**Fig. 2 C-E**). This result concurs with clinical observations about load induced sarcomere disarray(20, 35).

**Figure 4.**
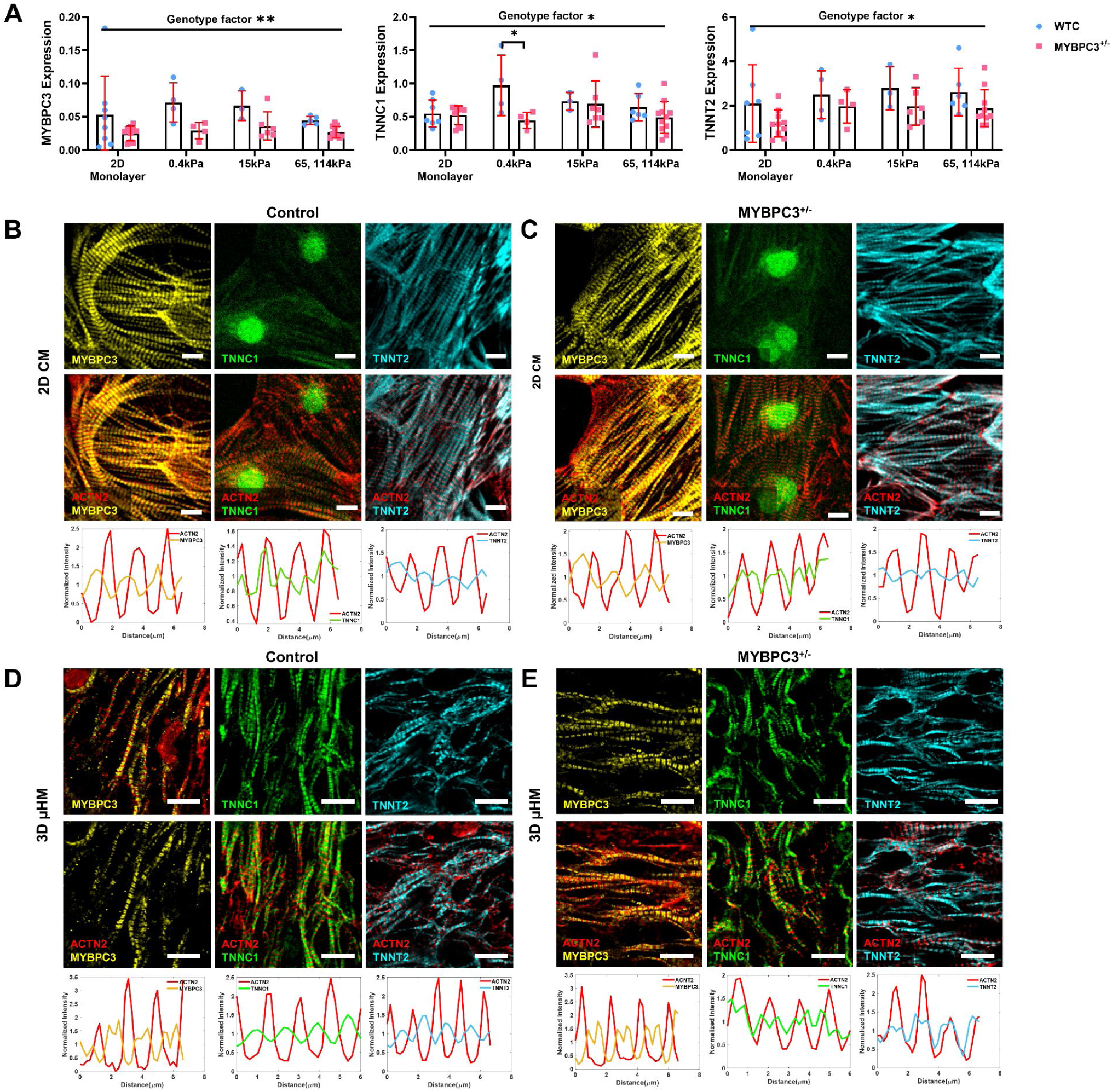
Mechanical stiffness trigger structural maturation of troponin C. (A) MYBPC3^+/-^ genotype has significantly less MYBPC3, troponin T and C expression compared to control. (B-C) Similar MYBPC3, TNNC1, and TNNT2 structural was seen between control and MYBPC3^+/-^ purified cardiomyocytes. Both MYBPC3 and TNNT2 have striated structure, however, troponin C is filamentous with localization at cellular nuclei (middle column). Targeted protein was merged with ACTN2, representative sarcomere traces were shown below each merged image, red indicates ACTN intensity, blue, green, cyan indicate MYBPC3, TNNC1, and TNNT2 respectively (D-E) Representative immunostaining for µHM formed on 15 kPa substrates. Compared to 2D sarcomere are widened in 3D, similar MYBPC3 localization was seen between the genotypes. Strikingly, we observed striated TNNC1 protein, yet both TNNC1 and TNNT2 protein have different localization in reference to ACTN2 compared to control tissues.

To test whether the structural abnormalities we observed were linked to differential adrenergic tone, as suggested by previous HCM studies, we stressed the µHMs with beta-adrenergic stimulation(37–39). Acute treatment with isoproterenol revealed a significant increase in spontaneous beat rate and calcium intake for both control and MYBPC3^+/-^ tissues (**Fig. 2 G-H**). MYBPC3^+/-^ tissues exhibited the same appropriate inotropic response to high dose isoproterenol (1 μM) as isogenic control tissues (**Fig. 2 H, SI Figure 4)**. These results suggest a preserved adrenergic reserve in the MYBPC3^+/-^ μHM. Together with evidence for cellular hypertrophy in the absence of gross sarcomere disarray, our model appears to reflect an early stage HCM phenotype(40, 41).

### Disturbed contractile kinetics and energetics in MYBPC3^+/-^ µHM

Clinical observations and mouse models of MYBPC3-linked HCM have suggested impairment of both diastolic relaxation and systolic contraction (42–45). Here, we observed that μHM with MYBPC3^+/-^ deficiency exhibited slower contraction kinetics compared to isogenic controls (**Fig. 3 A, C**). During spontaneous contraction, a hypercontractility phenotype was observed in MYBPC3^+/-^ μHM when substrate stiffness approached fibrotic levels (**Fig. 3 B**), consistent with the clinical phenotype of HCM, our own prior studies on µHM, and work by others on MYBPC3 mutant engineered heart muscle beating at low frequencies (≤1 Hz)(4, 22). However, analysis of contractile power curves during spontaneous beating revealed that the slower contraction kinetics of MYBPC3^+/-^ were associated with significantly more mechanical energy consumption for each contraction, but only when the tissues worked against substrates with physiological stiffness (15 kPa) (**Fig. 3 E, F**). Moreover, MYBPC3^+/-^

**Figure 3.**
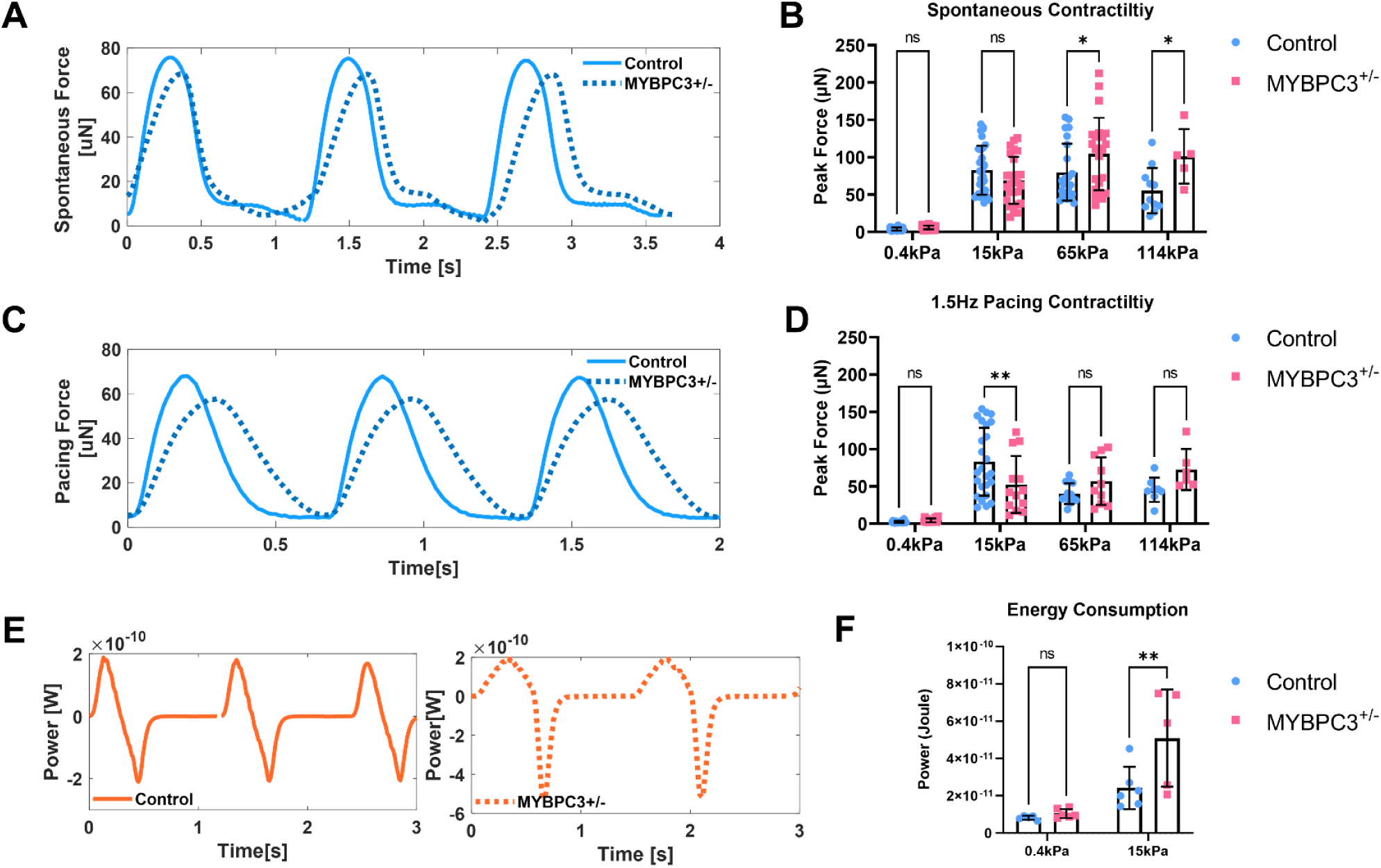
MYBPC3^+/-^ iPSC-µHM is associated with slower contraction kinetics together with impaired energy consumption. (A) Representative spontaneous force curves for both control and MYBPC3^+/-^ iPSC-µHM at 15 kPa, slower force rising phase was seen in MYBPC3^+/-^ iPSC-µHM. (B) During spontaneous contraction, a hypercontractility phenotype was seen in MYBPC3 tissues, especially at stiffer conditions. (C) Representative pacing force curves for both control and MYBPC3^+/-^ iPSC-µHM at 15 kPa, slower force kinetics with minimal resting phase was seen in MYBPC3^+/-^ iPSC-µHM. (D) At 1.5 Hz pacing, MYBPC3^+/-^ iPSC-µHM is hypocontracile at 15 kPa. (E) Representative power curve for both control and MYBPC3^+/-^ iPSC-µHM, a phenotype of non-uniform power distribution was seen in MYBPC3^+/-^ iPSC-µHM. (F) Integrated energy consumption indicated MYBPC3^+/-^ iPSC-µHM consume more power compared with control at 15 kPa. * and ** indicate *p* value less than 0.05 and 0.01. Error bar: *SD*.

μHM exhibited asymmetric power generation due to the slower rising phase of the contraction (**Fig. 3 E**). Likely because of their slower contraction kinetics, MYBPC3^+/-^ μHM exhibited significantly reduced contractility when field paced at a moderately high beat rate (1.5 Hz) and were *hypocontractile* on 15 kPa substrates (**Fig. 3 D**). These observations are consistent with prior animal and human iPSC-cardiomyocyte studies, where slower contraction kinetics in HCM cardiomyocytes, thought to be due to sarcomere abnormalities and disrupted super relaxed state, have been described(26, 46, 47).

### Differential troponin localization associated with MYBPC3^+/-^ µHM

Despite few differences observed in cardiomyocyte α-actinin organization (**Fig. 2**) or changes in transcription of ACTN2 (**SI Figure. 5**), we discovered that, along with the expected genetically encoded reduction in transcription of MYBPC3, transcript levels for troponin C (TNNC1), and cardiac troponin T (TNNT2) were also reduced substantially in MYBPC3^+/-^ tissues (**Fig. 4 A)**. To determine whether these RNA changes might be correlated to sarcomeric organization, we examined protein structure by immunostaining iPSC-cardiomyocytes in monoculture along with longitudinal cryosections of μHM. For day 30 purified iPSC-cardiomyocytes cultured in 2D, we observed a clear striated structure of ACTN2, TNNT2 and MYBPC3. In contrast, TNNC1 exhibited filamentous and nucleus localization in 2D iPSC-CM of both genotypes. This was reflected strongly in representative line traces of the sarcomere proteins, indicating significantly less periodicity of TNNC1 compared to other sarcomere proteins (**Fig. 4 B, C**).

In contrast to 2D cardiomyocyte cultures, cardiomyocytes within cryosections of control 3D μHM formed on physiological 15 kPa substrates showed more organized TNNC1 structure, with appropriate sarcomeric localization (**Fig. 4 D, E**). Moreover, within μHM of both genotypes, individual sarcomeres appeared to be wider than sarcomeres from 2D iPSC-CM (**Fig. 4 B-E**). Interestingly, in MYPBC3^+/-^ μHM, while MYBPC3 itself appeared to have appropriate localization, both TNNC1 and TNNT2 appear to be mis-localized. In isogenic control tissues, we observed physiological alternating localization of troponins with sarcomeric α-Actinin containing Z-disks, exemplified most readily in line-scans of the fluorescence micrographs. In contrast, within MYBPC3^+/-^ μHM, both troponin proteins exhibited potentially pathologic overlap with the Z-disk (**Fig. 4 D, E**).

### Mechanical resistance exacerbates calcium handling dysfunction in MYBPC3^+/-^ µHM

Calcium plays a critical role in cardiac contraction and relaxation; dysregulation in calcium can lead to abnormal contraction, arrhythmias, and structural changes in HCM(48). Thus, we assessed both action (49, 50) potential waveforms and calcium handling of the μHM (**Fig. 5 A, B**). In 2D iPSC-cardiomyocytes (differentiation day 30), we observed similar Ca^2+^ transients and action potential kinetics in cells of both genotypes, although MYBPC3^+/-^ iPSC-CM exhibited higher cytosolic calcium amplitude (ΔF/F_0_; *p* < 10^-4^, **Fig. 5 C**). However, culturing cardiomyocytes in μHM unveiled marked Ca^2+^ handling abnormalities in MYBPC3^+/-^ cells (**Fig. 5 B, E, F**). Most striking was a prolonged systolic Ca^2+^ plateau phase (**Fig. 5 B, D**) that was comparable to clinical observations of impaired myocardial relaxation, especially under chronic pressure overload conditions(25, 49). Abnormalities of Ca^2+^ dynamics in MYBPC3^+/-^ μHM, especially the prolonged systolic plateau, were exacerbated when μHM substrate stiffness exceeded 0.4 kPa (**Fig. 5 D**). Nevertheless, despite significant prolongation in Ca^2+^ transient, there were less profound differences in action potential waveform morphology (**Fig. 5 A, E, F**). These findings suggest that the primary electrophysiological difference between the early-stage HCM-prone MYBPC3^+/-^ tissues and isogenic controls is in Ca^2+^ handling, as opposed to a global electrophysiology change.

**Figure 5.**
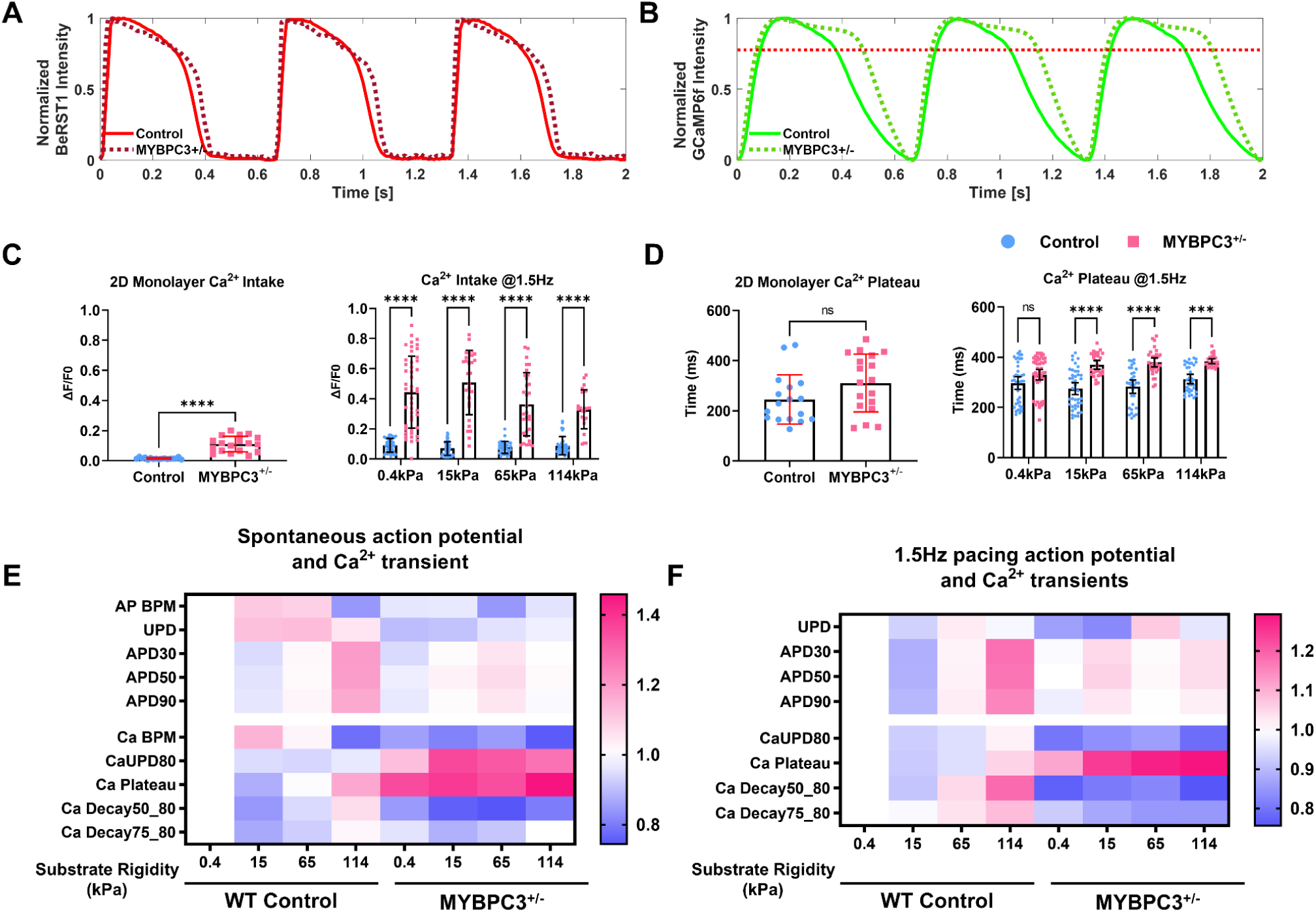
Mechanical stiffness trigger Ca^2+^ mishandling in MYBPC3^+/-^ iPSC-µHM. (A-B) Representative action potential and calcium traces for both control and MYBPC3^+/-^ iPSC-µHM at 15 kPa condition. Significantly prolonged Ca^2+^ during the peak was observed for MYBPC3^+/-^ iPSC-µHM. (C) Compared to 2D monolayer, iPSC-µHM has more Ca^2+^ intake, further, MYBPC3^+/-^ tissue has significantly higher Ca^2+^ intake compared to control. (D) 2D monolayer has similar Ca^2+^ kinetics compared to control. However, when cells formed in 3D tissue, a significantly longer Ca^2+^ plateau was observed at elevated substrate stiffnesses. (E-F) Heatmap of action potential and Ca^2+^ transient during spontaneous contraction (E) and 1.5 Hz paced contraction. Control tissue has prolonged action potential and Ca^2+^ transient at stiffness, while 15 kPa has the shortest action potential and Ca^2+^ transient. MYBPC3^+/-^ tissues always have significantly longer Ca^2+^ plateau compared to control, while MYBPC3^+/-^ tissues are less responsive to mechanical stiffness. *** and **** indicate p value less than 0.001 and 0.0001, error bar: *SD*.

### Pharmacological studies indicate L-type Ca^2+^ channel dysfunction in MYBPC3^+/-^ µHM

The prolonged Ca^2+^ plateau phase in MYBPC3^+/-^ tissues were not associated with transcriptional change of the *L*-type Ca^2+^ channel (CACNA1c, **Fig. 6 A**). A slight inhibition of SERCA transcript was observed in MYBPC3^+/-^ μHM (**Fig. 6 B**). SERCA is responsible for removal of >70% of cytoplasmic Ca^2+^(50), and SERCA transcript levels were observed to be reduced in cardiac biopsies(29, 51). To test whether Ca^2+^ mishandling was caused by SERCA inhibition, we administered a SERCA-specific inhibitor, thapsigargin(52). Thapsigargin treatment yielded a significant increase in Ca^2+^ upstroke duration that was nearly identical in magnitude for μHM of both genotypes (**Fig. 6 C**). Moreover, SERCA inhibition did not cause a prolonged Ca^2+^ plateau in control tissues (**SI Figure 6**). This drug also produced a trend toward hypercontractility in both genotypes, though the magnitude of the change was only significant in control tissues (**Fig. 6 D**). While the differential contractile response was notable, the fact that both control and MYBPC^+/-^ µHM exhibited similar Ca^2+^ handling changes in response to SERCA block strongly suggests against SERCA dysfunction being the primary mechanism for observed Ca^2+^ mishandling.

**Figure 6.**
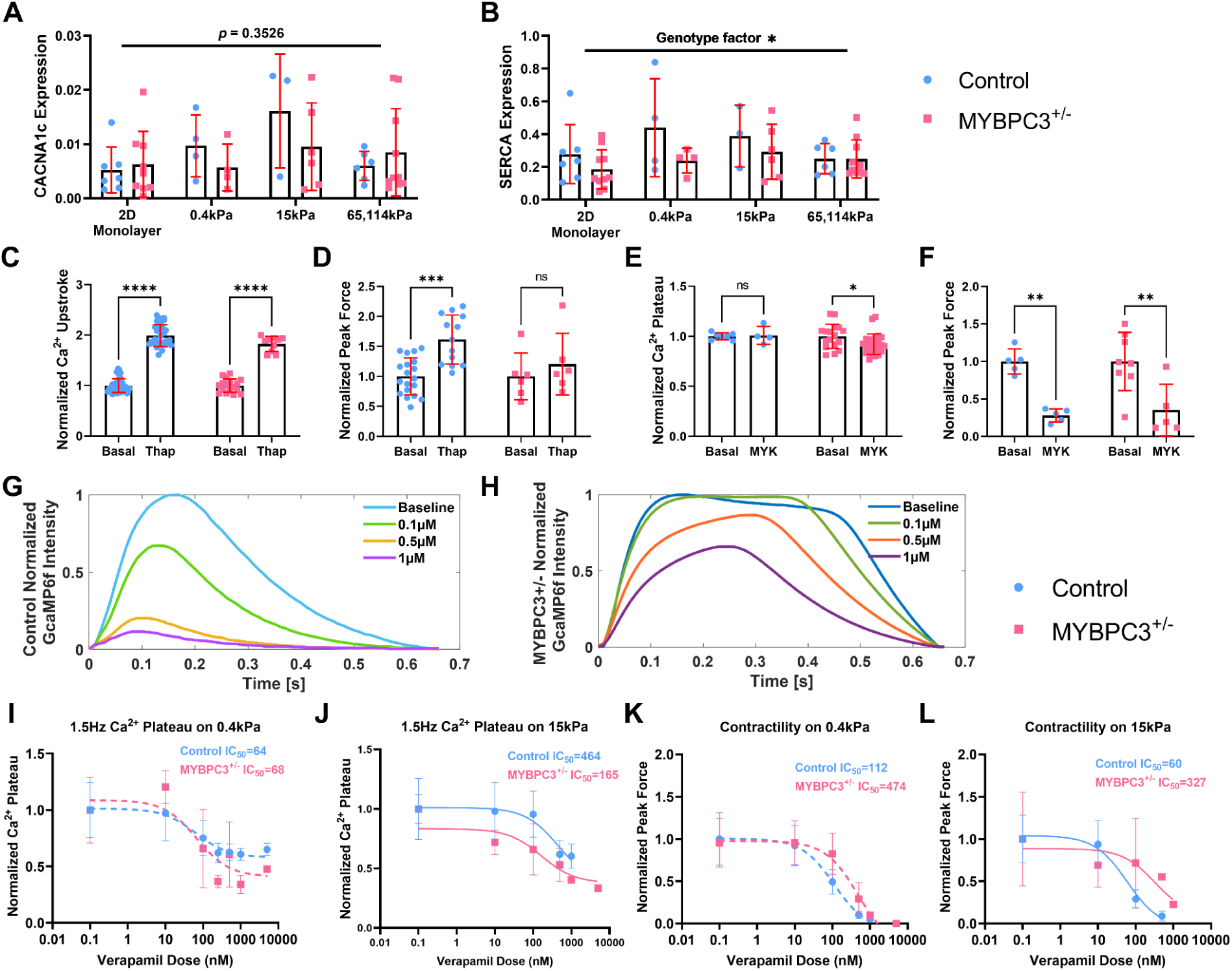
Pharmacological probes target specific Ca^2+^ mishandling mechanisms associated with MYBPC3. (A-B) Ca^2+^ regulation gene expression, similar CACNA1c and SERCA expression was seen between genotypes or stiffnesses. (C) Thapsigargin (Thap) has similar drug effects for both genotypes, significantly prolonged Ca^2+^ upstroke duration. (D) Thapsigargin resulted in higher contractility, especially for control tissues. (E) Mavacamten (MYK) did not change Ca^2+^ transient significantly, although there is a trend towards shortened Ca^2+^ plateau for MYBPC3^+/-^ tissues. (F) Significantly reduced contractility was seen for both genotypes with Mavacamten. (G-H) Control and MYBPC3^+/-^ µHM in response to verapamil. MYBPC3^+/-^ tissues showed more resistance to verapamil blocking. Higher doses of verapamil however reduce the Ca^2+^ plateau duration in MYBPC3^+/-^ µHM. (I-J) Ca^2+^ plateau duration in response to verapamil for both control and MYBPC3^+/-^ µHM, higher IC_50_ is required for higher mechanical resistance. (K-L) Contractility in response to verapamil for both control and MYBPC3^+/-^ µHM, higher IC_50_ is required for higher mechanical resistance. *, **, ***, and **** indicates *p* value less than 0.05, 0.01, 0.001 and 0.0001. Error bar: *SD*. n>4, 15 kPa tissues at 1.5 Hz was selected for comparison.

Dysfunction in Ca^2+^ handling in HCM has been suggested by some to originate directly from excessive Ca^2+^ buffering at myofilaments(53, 54). To test this possibility in our μHM, we acutely inhibited myosin using mavacamten(55). While others have reported that *chronic* mavacamten exposure can normalize Ca^2+^ handling in engineered heart muscle with HCM mutations(56), we observed only a very small change in the length of the systolic Ca^2+^ plateau in MYBPC3^+/-^ tissues with acute mavacamten dosing. However, it markedly reduces the contractility in both genotypes (**Fig. 6 E**). These results suggest that changes in myofilament activity do not acutely impact the overall Ca^2+^ transient. Consistent with our results, chronic mavacamten treatment was suggested to impact Ca^2+^ cycling through indirect means in prior work by others(57).

To investigate potential dysfunction of Ca^2+^ channels in MYBPC3^+/-^ μHM, we applied a Ca^2+^ channel blocker verapamil, which is widely prescribed for HCM patients (58). MYBPC3^+/-^ µHM exhibited markedly higher resistance to verapamil compared to control tissues: only very subtle changes in calcium handling were seen at 0.1 µM verapamil. In contrast, control tissues showed a significantly negative inotropic effect (**Fig. 6 G, H**). Unlike mavacamten, verapamil treatment significantly reduced the prolonged Ca^2+^ plateau phase observed in MYBPC3^+/-^ μHM. Importantly, MYBPC3^+/-^ tissues are even more resistant to verapamil at higher stiffnesses 15 kPa compared to 0.4 kPa (**Fig. 6 I, J**). This result suggests that although Ca^2+^ intake is increased in MYBPC3^+/-^ cardiomyocytes at baseline, mechanical stress provokes more exaggerated dysfunction of calcium handling. As with calcium handling, the impact of verapamil on force production in MYBPC3^+/-^ µHM was markedly less than the effect on control tissues, with the MYBPC3 deficient tissues exhibiting a much higher IC_50_: 474 vs 112 on 0.4 kPa condition and 327 vs 60 on 15 kPa conditions (**Fig 6 K, L**).

## Discussion

We developed a 3D physiological system with tunable mechanical resistance to promote cell maturation and to simulate coexistent hypertension, a known risk factor that worsens the prognosis of HCM patients(29). We optimized tissue formation to reduced batch-to-batch variation and expanded the substrate stiffness range from 0.4 to 114 kPa by simply using PDMS (**Fig. 1**)(4, 32). We characterized how the effects of mechanical stress can trigger hallmarks of HCM and recapitulate clinically relevant early-stage HCM phenotypes and drug responses(59).

We observed minimal differences in phenotypes between MYBPC3^+/-^ and isogenic control cardiomyocytes in 2D culture and in micro-tissues contracting against 0.4 kPa ultrasoft substrates (**Fig. 3**). Further, similar to 2D monolayers, low mechanical resistance 0.4 kPa caused less structural alignment of the sarcomere (**Fig. 2**), concordant with previous studies showing that proper mechanical stimuli increase engineered heart tissue maturity(8, 60, 61), and trigger proper tissue alignment(62–64). These observations further highlight the significance of environmental cues in modeling diseases with iPSC.

Surprisingly, while the MYPBC3 protein itself exhibited normal localization, we discovered a unique structural defect in troponin complex in MYBPC3^+/-^ tissues. This phenotype was masked in 2D culture and may be linked to structural maturation of the sarcomere triggered by 3D physiological environment. Within this tissue setting, we observed increased sarcomere width, and physiologic localization of TNNC1 in isogenic control tissues (**Fig. 2**, **4**). In purified 2D cardiac monolayers, we observed that TNNC1 was primarily localized to the nucleus rather than the sarcomere. Similar mis-localization of this protein has been reported by others in neonatal rat ventricular cardiomyocyte and iPSC-cardiomyocytes (65, 66). Our observation of defective localization of TNNC1 and TNNT2 in μHM mirrors clinical observations of HCM patients, where cardiac cell degeneration is associated with structural defects including an abortive form of sarcomerogenesis(67). The micro-defects of troponin complex may be an early structural deformation associated with MYBPC3 mutations, potentially relating the physiological phenotypes observed. For example, the troponin complex is strongly linked to heart muscle contraction and can cause Ca^2+^ sensitivity(68–70). Further, troponins have important implications in cardiomyopathy, with identified troponin mutations causing hypertrophic, dilated, and restrictive myopathies(48, 71, 72).

In addition to the unique troponin structural defects, we captured marked changes in the Ca^2+^ transient for the MYBPC3^+/-^ tissues, with significantly higher Ca^2+^ intake and a prolonged systolic Ca^2+^ plateau. Ca^2+^ mishandling has been reported in both human and mice HCM models, with potential mechanisms including dysfunction of Ca^2+^ channels, disrupted Ca^2+^ buffering, and stress induced Ca^2+^ sensitivity(26–29, 48, 73). While SERCA inhibition has been reported in obstructive HCM patient samples, we observed similar SERCA function between control and MYBPC3^+/-^ tissues, indicating SERCA may not be the primary cause of the prolonged Ca^2+^ plateau in our µHM model(29, 74).

Meanwhile, the hypothesis of altered myofilament Ca^2+^ buffering has been supported by studies with recombinant protein and isolated pig heart samples associated with thin filament proteins like troponin(28, 57). However, we and others observed that mavacamten, which shifts myosin into their super-relaxed state(45), has much less effect on intracellular Ca^2+^ compared to its effect on contraction inhibition. This argues against dramatic changes in calcium buffering at the myofilaments as being a direct cause in HCM-linked calcium abnormalities. Whether chronic depression of HCM-cardiomyocyte contractility with mavacamten can cause adaptive reduction of calcium buffering needs to be further investigated(48, 75, 76). We observed that aberrant Ca^2+^ handling is associated with a much slower contraction kinetics and impaired energy consumption in MYBPC3^+/-^ µHM.

Our observations of differential verapamil responses between control and MYBPC3^+/-^ μHM suggest an early and excessive activation of *L*-type calcium channel flux in MYBPC3^+/-^ µHM, potentially altering the intercellular Ca^2+^ homeostasis, and leading to impaired force generation kinetics and hypertrophy(26, 47). Notably, while MYBPC3^+/-^ μHM on 0.4 kPa substrates had similar verapamil response to controls, the genotype-linked difference was readily observed on substrates with physiologic stiffness. Consistent with our observations of hypertrophy (**Fig. 2**) and contractile energetics (**Fig 3**), we find that mechanical resistance to contractility - linked to substrate stiffness and *in vivo* linked to hypertension - plays a critical role in early-stage pathogenesis of MYBPC3-linked HCM(4, 77, 78).

In summary, we have developed a new μHM model that leverages substrates with tunable stiffness to investigate how non-genetic factors such as hypertension may be synergistic with genomic variants in triggering pathogenesis of HCM. Our simple PDMS based physiological model provides a useful tool to understand the pathogenesis of HCM. It will be helpful for the development and evaluation of novel pharmacological targets. Importantly, this system can be widely employed to explore other inherited cardiomyopathies and investigate combined effects of cardiac mechanical overload and genetic mutations.

## Materials and Methods

### Material characterization

To address our hypothesis that mechanics can initiate HCM pathogenesis *in vitro*, we formed µHM on PDMS substrates ranging in elastic modulus from 0.4 kPa up to 114 kPa (Fig. 1, **SI Figure. 1**) (17, 29, 30). Polydimethylsiloxane (PDMS) elastomer was used for making substrates and stencil molds. Elastic modulus of the material was characterized using a planar biaxial testing machine(32).

Ultrasoft 0.4 kPa substrates mimicking the embryonic heart microenvironment were made using sylgard 527 (5:4 ratio of component A to component B). Substrates with rigidity ranging from values near that of healthy adult human heart (15 kPa) to fibrosis-like (65kPa and 114kPa) substrates were made by mixing sylgard 184 into sylgard 527 with ratio of 1:40, 1:10 and 1:5(20, 30, 31).

“Dog bone” shaped stencil molds to control the shape of micro-heart muscles were made of sylgard 184 using 3D printed masters(79). Briefly, “dog bone” positive 3D prints were generated from FormLabs Clear Resin^TM^. To avoid toxicity from direct molding sylgard 184 off the 3D prints, 1.5% agar gels were cast directly off to make “dog bone” negative. Sylgard 184 was then molded off the agar gel to get a positive replica of the original 3D print. Finally, the PDMS replicas of the 3D prints were treated with fluorosilane before Sylgard 184 stencils were cast off these replica PDMS molds(32, 80). PDMS was cured against the replica mold under tight compression so that the “dog bone” shapes formed through-holes through the PDMS stencils.

### Device fabrication

μHM forming stencils were prepared on soft substrates based on our previously described methods(4, 32). Briefly, a thick PDMS base layer (∼ 200 µm thick) and a thinner PDMS/fluorescent bead layer (∼ 5 µm thick) were applied to glass coverslips by spin coating. Soft PDMS substrates were cured sequentially overnight at 60°C. Substrates were then covalently bound to the ECM protein fibronectin following plasma oxidation, amino-silane treatment, and glutaraldehyde modification(4). Following covalent bonding of fibronectin, the modified surface was further quenched with 2.5 w/v% ethanolamine (in dPBS). Pluronic treated sylgard 184 stencils were then absorbed onto the substrates with methanol. Finally, the device was disinfected by applying 70% ethanol overnight.

### Human induced pluripotent stem cell culture and differentiation

MYBPC3^+/-^ iPSC and isogenic controls of the Wild Type C background (WTC; Coriell GM2525) harboring the knock-in GCamP6f calcium indicator were cultured in Essential 8 media and were differentiated to cardiomyocytes using Wnt pathway modulators. Using this method spontaneous contraction could be observed within 10 days of starting differentiation by treating iPSC with CHIR99021 to initiate Wnt signaling (4, 32). On differentiation day 15, beating cardiomyocyte monolayers were singularized (0.25% trypsin) and replated onto Matrigel coated tissue culture plastic at 250k/cm^2^ for lactate purification as previously described(32). Cardiomyocytes purities were quantified using flow cytometry sorting for cardiac troponin T positive cells. At differentiation day 30, purified cardiomyocytes were used for making micro heart muscle as described previously(4).

### iPSC-µHM Formation and Optimization

Purified cardiomyocytes were gently dissociated using 0.25% trypsin EDTA (3-4 mins) and seeded into each “dog bone” shaped mold with 3 µL of 7 x 10^7^ cell/mL (210,000 cells). The cells were then incubated at 37°C with 5% CO_2_ for 2 hours before the entire culture well was filled with replating media (Dulbecco’s Modified Eagle Media, DMEM supplemented with 20% FBS, 150 μg/mL ascorbic acid and 10 μM Y27632). Tissue compaction and alignment was typically observed within 24 hours(4). At least 4 high quality tissue batches with cardiomyocytes purity of 85±10% were selected for study. Tissue quality was determined by tissue shape, visible cellular death, and maintenance of adhesion to the substrates (**SI Figure. 3)**. We used tissue gross shape as our first quality control metric because of our recent findings that overall μHM shape impacts physiology(80). Tissues that deviated significantly from the expected “dog bone” shape or were delaminated from the substrate were excluded. We observe the inclusion/exclusion rate are not dependent on genotypes or stiffnesses (**SI Figure. 3**). Altogether, these quality control metrics led to a more consistent and narrower range of μHM physiologic behaviors compared to what was previously possible(32).

### Gene Expression Analysis

Expression of sarcomere genes and calcium genes were evaluated with quantitative RT-PCR(4). Detailed gene primer sequence in supplemental. Cell or tissue lysis and RNA recovery were performed using RNAqueous®-Micro Kit (Thermo Scientific). At least 320 ng RNA per sample was synthesized for cDNA using Super Script III Reverse Transcriptase Kit (Thermo Scientific). Gene expression was performed using Applied Biosystems Step One Plus Light Cycler instrument using SYBR green master mix (Thermo Scientific).

### Immunostaining

The tissues were step fixed with 1%, 2%, 3% and 4% paraformaldehyde at 4°C, with each fixation step at least 1 hour(8). Then was transferred to 1% agar followed by OCT protection as described previously(32). The 8 µm cryosections were used for immunostaining. Sarcomere α-actinin, myosin binding protein C, troponin T, troponin C and wheat germ agglutinin (WGA, **SI Table 2**), together with cell nuclei were stained and scanned using Olympus confocal model (Fluoview FV1200, Tokyo, Japan). Tissue alignments were measured by using sarcomere alfa actinin staining, the angles between individual sarcomere to the tissue shaft were measured through custom built MATLAB code. Cell size was quantified by drawing individual cross-sectional areas of cell using ImageJ. Sarcomere profiles were plotted by selecting representative sarcomere regions in ImageJ and normalized by the mean intensity value of the specific sarcomere trace.

### High-Speed Fluorescent Imaging

Physiology of iPSC-μHM at day 10 (corresponding to day 40 of iPSC-CM differentiation) was analyzed using high speed imaging system (Nikon Eclipse Ts2R; Tokyo, Japan) equipped with a digital CMOS camera (Hamamatsu ORCA-Flash4.0 V2; Japan)(32). The imaging stage was set to a constant 37°C temperature using a thermal plate (Tokai Hit; Shizuoka, Japan)(4). The field pacing of tissue (1.5 Hz, 20msec Bipolar pulses, <20V) during imaging was achieved using pacing graphite electrodes (MyoPacer, Ion Optix, USA).

### Contractility characterization

Traction force microscopy (TFM) was used to quantify tissue contractility. Tissue peak force was calculated through using substrate elasticity and measuring green fluorescence beads displacement was characterized via open-source software as described previously(4)(81). For TFM, we assumed the Poisson’s ratio of PDMS was 0.49, detailed parameters for TFM analysis are based on prior work(32, 81, 82).

### Mechanical energy consumption calculation

Instantaneous contractile power was calculated at each imaging timepoint by multiplying force and velocity obtained from TFM calculations. These curves were used to calculate energy consumed during contraction and relaxation(81). Briefly, positive, and negative regions of the power curve from each contraction cycle, were integrated by calculating the area under the curve, positive energy consumptions were used for comparison between μHM of each genotype.

### Action potential and calcium measurements

Membrane potential was measured by labeling tissue overnight with 1µM BeRST-1 voltage sensitive dye in phenol red free RPMI-C(83). Action potential waveforms were captured by imaging in the Cy5 channel. Calcium transients were captured by imaging endogenously expressed GcaMP6f in the GFP channel(4). Videos was analyzed via custom build code; intensity profile and kinetics parameters were automatically extracted(84). For calcium transient dynamics, initial baseline to 80% of the intensity was measured as UPD80, plateau region of more than 80% intensity was measured, decay times from end of the plateau to 50 and 75% decay of intensity was characterized.

### Pharmacology studies

The potential mechanism for pathologic calcium handling in HCM µHM was investigated using pharmacological probes. Specifically, isoproterenol (0-1 µM) L-type Ca^2+^ blocker verapamil (0-1 µM), SERCA inhibitor thapsigargin (0, 5 µM) and myosin inhibitor mavacamten (0, 0.5 µM)(52, 55, 85). Drugs were first diluted in dimethyl sulfoxide, then serial diluted in RPMI/C media with supplements. Acute drug administration was performed by at least 45 minutes incubation at 37°C with 5% CO_2_ prior imaging.

### Statistics

GraphPad 9.3.1 was used for statistics. One-way or two-way analysis of variance (ANOVA) was used followed by multiple comparison test between groups was performed using Holm Sidak’s method. The *p* value less than 0.05 was considered statistically significance.

## Supporting information

Guo et al HCM mechanism 2023 SI

## Acknowledgments

We thank Drs. Guy M Genin, Sharon Cresci, Jonathan Moreno, Michael Vahey, Lori Setton, Jianjun Guan for helpful discussions and use of the equipment. We thank Dr. Evan Miller for providing BeRST1 dye. This work was supported by the Departments of Biomedical Engineering and Mechanical Engineering and Material Science at Washington University in St. Louis, the American Heart Association (19CDA34730016 to NH, predoctoral fellowship 828938 to JG, TPA 970198 to NH) and the National Institutes of Health (R01 HL159094 to NH).

## References

1. M. Lorenzini, et al., Penetrance of Hypertrophic Cardiomyopathy in Sarcomere Protein Mutation Carriers. J Am Coll Cardiol 76, 550–559 (2020).

2. E. J. Rowin, B. J. Maron, M. S. Maron, The Hypertrophic Cardiomyopathy Phenotype Viewed Through the Prism of Multimodality Imaging: Clinical and Etiologic Implications. JACC Cardiovasc Imaging 13, 2002–2016 (2020).

3. G. G. Repetti, et al., Discordant clinical features of identical hypertrophic cardiomyopathy twins 10.1073/pnas.2021717118/-/DCSupplemental.

4. J. Guo, et al., Interplay of Genotype and Substrate Stiffness in Driving the Hypertrophic Cardiomyopathy Phenotype in iPSC-Micro-Heart Muscle Arrays. Cell Mol Bioeng 14, 409–425 (2021).

5. Z. Ma, et al., Contractile deficits in engineered cardiac microtissues as a result of MYBPC3 deficiency and mechanical overload. Nat Biomed Eng 2, 955–967 (2018).

6. D. Barefield, et al., Haploinsufficiency of MYBPC3 exacerbates the development of hypertrophic cardiomyopathy in heterozygous mice. J Mol Cell Cardiol 79, 234–243 (2015).

7. S. R. Ommen, et al., 2020 AHA/ACC Guideline for the Diagnosis and Treatment of Patients With Hypertrophic Cardiomyopathy: A Report of the American College of Cardiology/American Heart Association Joint Committee on Clinical Practice Guidelines. (2020) 10.1161/CIR.0000000000000937.

8. K. Ronaldson-Bouchard, et al., Advanced maturation of human cardiac tissue grown from pluripotent stem cells. Nature 556, 239–243 (2018).

9. F. Weinberger, I. Mannhardt, T. Eschenhagen, Engineering Cardiac Muscle Tissue: A Maturating Field of Research. Circ Res 120, 1487–1500 (2017).

10. P. J. M. Wijnker, J. van der Velden, Mutation-specific pathology and treatment of hypertrophic cardiomyopathy in patients, mouse models and human engineered heart tissue. Biochim Biophys Acta Mol Basis Dis 1866 (2020).

11. C. Liu, A. Oikonomopoulos, N. Sayed, J. C. Wu, Modeling human diseases with induced pluripotent stem cells: From 2D to 3D and beyond. Development (Cambridge) 145, 1–6 (2018).

12. B. J. Maron, M. S. Maron, Hypertrophic cardiomyopathy. The Lancet 381, 242–255 (2013).

13. H. J. Chang, C. Lynm, R. M. Glass, Hypertrophic cardiomyopathy. JAMA - Journal of the American Medical Association 302, 1720 (2009).

14. S. Marston, O. Copeland, K. Gehmlich, S. Schlossarek, L. Carrrier, How do MYBPC3 mutations cause hypertrophic cardiomyopathy? J Muscle Res Cell Motil 33, 75–80 (2012).

15. 15. A. S. Helms, et al., Effects of MYBPC3 loss-of-function mutations preceding hypertrophic cardiomyopathy. *JCI Insight* 5, e133782 (2020).

16. C. Y. Ho, et al., Valsartan in early-stage hypertrophic cardiomyopathy: a randomized phase 2 trial. Nat Med 27, 1818–1824 (2021).

17. C. Y. Ho, et al., Diltiazem treatment for pre-Clinical hypertrophic cardiomyopathy sarcomereMutation carriers: A pilot randomized trial to modify disease expression. JACC Heart Fail 3, 180–188 (2015).

18. N. Frey, M. Luedde, H. A. Katus, Mechanisms of disease: Hypertrophic cardiomyopathy. Nat Rev Cardiol 9, 91–100 (2012).

19. B. A. Maron, et al., What Causes Hypertrophic Cardiomyopathy? American Journal of Cardiology 179, 74–82 (2022).

20. A. M. Varnava, P. M. Elliott, S. Sharma, W. J. McKenna, M. J. Davies, Hypertrophic cardiomyopathy: The interrelation of disarray, fibrosis and small vessel disease. Heart 84, 476– 482 (2000).

21. J. C. Tardiff, et al., Cardiac troponin T mutations result in allele-specific phenotypes in a mouse model for hypertrophic cardiomyopathy. Journal of Clinical Investigation 104, 469–481 (1999).

22. R. Cohn, et al., A Contraction Stress Model of Hypertrophic Cardiomyopathy due to Sarcomere Mutations. Stem Cell Reports 12, 71–83 (2019).

23. A. S. vander Roest, et al., Hypertrophic cardiomyopathy β-cardiac myosin mutation (P710R) leads to hypercontractility by disrupting super relaxed state. PNAS 118 (2021).

24. M. A. Trembley, et al., Mechanosensitive gene regulation by myocardin-related transcription factors is required for cardiomyocyte integrity in load-induced ventricular hypertrophy. Circulation 138, 1864–1878 (2018).

25. R. Coppini, et al., Late sodium current inhibition reverses electromechanical dysfunction in human hypertrophic cardiomyopathy. Circulation 127, 575–584 (2013).

26. R. Coppini, C. Ferrantini, A. Mugelli, C. Poggesi, E. Cerbai, Altered Ca2+ and Na+ homeostasis in human hypertrophic cardiomyopathy: Implications for arrhythmogenesis. Front Physiol 9, 1–16 (2018).

27. F. Lan, et al., Abnormal calcium handling properties underlie familial hypertrophic cardiomyopathy pathology in patient-specific induced pluripotent stem cells. Cell Stem Cell 12, 101–113 (2013).

28. P. Robinson, et al., Hypertrophic cardiomyopathy mutations increase myofilament Ca2 buffering, alter intracellular Ca2 handling, and stimulate Ca2-dependent signaling. Journal of Biological Chemistry 293, 10487–10499 (2018).

29. A. S. Helms, et al., Genotype-Dependent and -Independent Calcium Signaling Dysregulation in Human Hypertrophic Cardiomyopathy. Circulation 134, 1738–1748 (2016).

30. A. J. Engler, et al., Embryonic cardiomyocytes beat best on a matrix with heart-like elasticity: Scar-like rigidity inhibits beating. J Cell Sci 121, 3794–3802 (2008).

31. B. Hinz, Tissue Stiffness, Latent TGF-β1 Activation, and Mechanical Signal Transduction: Implications for the Pathogenesis and Treatment of Fibrosis. Curr Rheumatol Rep 11, 120–126 (2009).

32. J. Guo, et al., Elastomer-Grafted iPSC-Derived Micro Heart Muscles to Investigate Effects of Mechanical Loading on Physiology. ACS Biomater Sci Eng (2020) 10.1021/acsbiomaterials.0c00318.

33. C. H. Conrad, et al., Myocardial Fibrosis and Stiffness With Hypertrophy and Heart Failure in the Spontaneously Hypertensive Rat. Circulation 91, 161–170 (1995).

34. M. Yildiz, et al., Left ventricular hypertrophy and hypertension. Prog Cardiovasc Dis 63, 10–21 (2020).

35. M. A. Trembley, et al., Mechanosensitive gene regulation by myocardin-related transcription factors is required for cardiomyocyte integrity in load-induced ventricular hypertrophy. Circulation 138, 1864–1878 (2018).

36. D. Zhang, et al., Tissue-engineered cardiac patch for advanced functional maturation of human ESC-derived cardiomyocytes. Biomaterials 34, 5813–5820 (2013).

37. R. Doste, R. Coppini, A. Bueno-Orovio, Remodelling of potassium currents underlies arrhythmic action potential prolongation under beta-adrenergic stimulation in hypertrophic cardiomyopathy. J Mol Cell Cardiol 172, 120–131 (2022).

38. N. Sun, et al., “Patient-Specific Induced Pluripotent Stem Cells as a Model for Familial Dilated Cardiomyopathy.”

39. A. Elesber, et al., Utility of Isoproterenol to Provoke Outflow Tract Gradients in Patients With Hypertrophic Cardiomyopathy. American Journal of Cardiology 101, 516–520 (2008).

40. A. Elesber, et al., Utility of Isoproterenol to Provoke Outflow Tract Gradients in Patients With Hypertrophic Cardiomyopathy. American Journal of Cardiology 101, 516–520 (2008).

41. R. Doste, R. Coppini, A. Bueno-Orovio, Remodelling of potassium currents underlies arrhythmic action potential prolongation under beta-adrenergic stimulation in hypertrophic cardiomyopathy. J Mol Cell Cardiol 172, 120–131 (2022).

42. S. J. van Dijk, et al., Contractile Dysfunction Irrespective of the Mutant Protein in Human Hypertrophic Cardiomyopathy With Normal Systolic Function (2011) 10.1161/CIRCHEARTFAILURE.111.963702/-/DC1.

43. G. Mearini, et al., Mybpc3 gene therapy for neonatal cardiomyopathy enables long-term disease prevention in mice. Nat Commun 5 (2014).

44. M. J. Oliva-Sandoval, et al., Insights into genotype-phenotype correlation in hypertrophic cardiomyopathy. Findings from 18 Spanish families with a single mutation in MYBPC3. Heart 96, 1980–1984 (2010).

45. C. N. Toepfer, et al., Hypertrophic cardiomyopathy mutations in MYBPC3 dysregulate myosin. Sci Transl Med 11 (2019).

46. D. B. S. V. R. Designed Research; A, K. M. R. S. V. R. Performed Research; A, Hypertrophic cardiomyopathy β-cardiac myosin mutation (P710R) leads to hypercontractility by disrupting super relaxed state. 118 (2021).

47. C. Ferrantini, et al., Pathogenesis of hypertrophic cardiomyopathy is mutation rather than disease specific: A comparison of the cardiac troponin T E163R and R92Q mouse models. J Am Heart Assoc 6 (2017).

48. G. L. Smith, D. A. Eisner, Calcium Buffering in the Heart in Health and Disease. Circulation 139, 2358–2371 (2019).

49. B. H. Lorell, W. Grossman, Cardiac hypertrophy: The consequences for diastol. J Am Coll Cardiol 9, 1189–1193 (1987).

50. D. H. MacLennan, E. G. Kranias, Phospholamban: A crucial regulator of cardiac contractility. Nat Rev Mol Cell Biol 4, 566–577 (2003).

51. R. Coppini, et al., Late sodium current inhibition reverses electromechanical dysfunction in human hypertrophic cardiomyopathy. Circulation 127, 575–584 (2013).

52. J. Lyttonsg, M. Westlins, M. R. Hanleyll, Thapsigargin Inhibits the Sarcoplasmic or Endoplasmic Reticulum Ca-ATPase Family of Calcium Pumps*. THE JOURNAL OF BIOLOGICAL CHEMISTRY 266, 17067–17071 (1991).

53. C. Semsarian, et al., The L-type calcium channel inhibitor diltiazem prevents cardiomyopathy in a mouse model. Journal of Clinical Investigation 109, 1013–1020 (2002).

54. A. J. Sparrow, et al., Measurement of Myofilament-Localized Calcium Dynamics in Adult Cardiomyocytes and the Effect of Hypertrophic Cardiomyopathy Mutations. Circ Res 124, 1228– 1239 (2019).

55. E. M. Green, et al., Heart disease: A small-molecule inhibitor of sarcomere contractility suppresses hypertrophic cardiomyopathy in mice. Science *(1979)* 351, 617–621 (2016).

56. L. R. Sewanan, et al., Loss of crossbridge inhibition drives pathological cardiac hypertrophy in patients harboring the tpm1 e192k mutation. Journal of General Physiology 153 (2021).

57. A. J. Sparrow, H. Watkins, M. J. Daniels, C. Redwood, P. Robinson, Mavacamten rescues increased myofilament calcium sensitivity and dysregulation of Ca 2 flux caused by thin filament hypertrophic cardiomyopathy mutations. Am J Physiol Heart Circ Physiol 318, 715–722 (2020).

58. R. O Bonow, et al., Effects of Verapamil on Left Ventricular Systolic Function and Diastolic Filling in Patients with Hypertrophic Cardiomyopathy. 787–796 (1987).

59. S. J. Lehman, C. Crocini, L. A. Leinwand, Targeting the sarcomere in inherited cardiomyopathies. Nat Rev Cardiol 19, 353–363 (2022).

60. A. Leonard, et al., Afterload promotes maturation of human induced pluripotent stem cell derived cardiomyocytes in engineered heart tissues. J Mol Cell Cardiol 118, 147–158 (2018).

61. D. Zhang, et al., Tissue-engineered cardiac patch for advanced functional maturation of human ESC-derived cardiomyocytes. Biomaterials 34, 5813–5820 (2013).

62. T. Eschenhagen, et al., Three-dimensional reconstitution of embryonic cardiomyocytes in a collagen matrix: A new heart muscle model system. FASEB Journal 11, 683–694 (1997).

63. S. S. Nunes, et al., Biowire: A platform for maturation of human pluripotent stem cell-derived cardiomyocytes. Nat Methods 10, 781–787 (2013).

64. W. Bian, N. Badie, H. D. Himel, N. Bursac, Robust T-tubulation and maturation of cardiomyocytes using tissue-engineered epicardial mimetics. Biomaterials 35, 3819–3828 (2014).

65. F. Z. Asumda, P. B. Chase, Nuclear cardiac troponin and tropomyosin are expressed early in cardiac differentiation of rat mesenchymal stem cells. Differentiation 83, 106–115 (2012).

66. H. Wu, et al., Epigenetic Regulation of Phosphodiesterases 2A and 3A Underlies Compromised β-Adrenergic Signaling in an iPSC Model of Dilated Cardiomyopathy. Cell Stem Cell 17, 89–100 (2015).

67. B. J. Maron, V. J. Ferrans, W. C. Roberts, “Ultrastructural Features of Degenerated Cardiac Muscle Cells in Patients With Cardiac Hypertrophy” (1975).

68. M. X. Li, P. M. Hwang, Structure and function of cardiac troponin C (TNNC1): Implications for heart failure, cardiomyopathies, and troponin modulating drugs. Gene 571, 153–166 (2015).

69. I. A. Katrukha, Human cardiac troponin complex. structure and functions. Biochemistry (Moscow) 78, 1447–1465 (2013).

70. J. K. Siddiqui, et al., Myofilament Calcium Sensitivity: Consequences of the effective concentration of troponin I. Front Physiol 7 (2016).

71. S. Dewan, K. J. McCabe, M. Regnier, A. D. McCulloch, S. Lindert, Molecular Effects of cTnC DCM Mutations on Calcium Sensitivity and Myofilament Activation - An Integrated Multiscale Modeling Study. Journal of Physical Chemistry B 120, 8264–8275 (2016).

72. D. Lin, A. Bobkova, E. Homsher, L. S. Tobacman, “Functional analyses of troponin T mutations that cause hypertrophic cardiomyopathy: Insights into disease pathogenesis and troponin function” (1998).

73. H. E. Cingolani, N. G. Pérez, O. H. Cingolani, I. L. Ennis, The Anrep effect: 100 years later. Am J Physiol Heart Circ Physiol 304, 175–182 (2013).

74. K. Toischer, et al., Elevated afterload, neuroendocrine stimulation, and human heart failure increase BNP levels and inhibit preload-dependent SERCA upregulation. Circ Heart Fail 1, 265– 271 (2008).

75. T. Sorsa, P. Pollesello, R. J. Solaro, “The contractile apparatus as a target for drugs against heart failure: Interaction of levosimendan, a calcium sensitiser, with cardiac troponin c” (Kluwer Academic Publishers, 2004).

76. L. R. Sewanan, et al., Loss of crossbridge inhibition drives pathological cardiac hypertrophy in patients harboring the tpm1 e192k mutation. Journal of General Physiology 153 (2021).

77. A. F. Leite-Moreira, J. Correia-Pinto, T. C. Gillebert, Afterload induced changes in myocardial relaxation: A mechanism for diastolic dysfunction. Cardiovasc Res 43, 344–353 (1999).

78. R. Truitt, et al., Increased Afterload Augments Sunitinib-Induced Cardiotoxicity in an Engineered Cardiac Microtissue Model. JACC Basic Transl Sci 3, 265–276 (2018).

79. D. W. Simmons, D. R. Schuftan, G. Ramahdita, N. Huebsch, Hydrogel-Assisted Double Molding Enables Rapid Replication of Stereolithographic 3D Prints for Engineered Tissue Design. ACS Appl Mater Interfaces 15, 25313–25323 (2023).

80. D. W. Simmons, et al., Hydrogel Assisted Double Molding of 3D-Print Enables Prestress Regulation of Micro-Heart Muscle Physiology 10.1101/2022.07.23.501265.

81. A. J. S. Ribeiro, et al., Multi-imaging method to assay the contractile mechanical output of micropatterned human iPSC-derived cardiac myocytes. Circ Res 120, 1572–1583 (2017).

82. S. Dogru, B. Aksoy, H. Bayraktar, B. E. Alaca, Poisson’s ratio of PDMS thin films. Polym Test 69, 375–384 (2018).

83. Y. L. Huang, A. S. Walker, E. W. Miller, A Photostable Silicon Rhodamine Platform for Optical Voltage Sensing. J Am Chem Soc 137, 10767–10776 (2015).

84. K. Oguntuyo, et al., Robust, Automated Analysis of Electrophysiology in Induced Pluripotent Stem Cell-Derived Micro-Heart Muscle for Drug Toxicity. Tissue Eng Part C Methods (2022) 10.1089/ten.tec.2022.0053.

85. N. Huebsch, et al., Miniaturized iPS-Cell-Derived Cardiac Muscles for Physiologically Relevant Drug Response Analyses. Sci Rep 6, 1–12 (2016).

