## Supplementary material for "Mechanical Resistance to Micro-Heart Tissue Contractility unveils early Structural and Functional Pathology in iPSC Models of Hypertrophic Cardiomyopathy": Guo et al HCM mechanism 2023 SI: Guo et al HCM Mechanism 2023 SI.pdf

**Supplemental Information for**
**Mechanical Loading unveils Calcium Handling Dysfunction in iPSC micro-heart muscle harboring Hypertrophic Cardiomyopathy Mutations**

| Gene Primer | Forward | Reverse | Vendor |
|---|---|---|---|
| GAPDH | CTCTGCTCCTCCTGTTCGAC | TTAAAAGCAGCCCTGGTGAC | IDT |
| ACTN2 | GCTTCTACCACGCTTTTGCG | CATTCCAAAAGCTCACTCGCT | IDT |
| TNNT2 | TTCACCAAAGATCTGCTCCTCGCT | TTATTACTGGTGTGGAGTGGGTGTGG | IDT |
| MYBPC3 | GGCATGCTAAAGAGGCTCAA | TCTTGTGGCCTTTGCTCAC | IDT |
| SERCA | ACCCACATTCGAGTTGGAAG | CCAACGAAGGTCAGATTGGT | IDT |
| CACNA1C | GGAGAGTTTTCCAAAGAGAG | TTTGAGATCCTCTTCTAGCTG | IDT |
| TNNC1 | ATGAGCTGAAGATAATGCTG | AACTCCAGGAACTCATCATAG | Sigma |

**Supplemental Table 1: Primers for Quantitative RT-PCR.**

| Protein Antigen | Clone, Catalog# and vendor | Host species and reactivity | Concentration or fold dilution used and application |
|---|---|---|---|
| Cardiac α-actinin (ACTN2) | EA-53, A7811, Sigma | Mouse/IgG1, monoclonal | 1:1000 IHC, 1:2500 Western |
| Cardiac α-actinin (ACTN2) | 1E11, ZRB1169, Sigma | Rabbit/IgG, recombinant monoclonal | 1:100 IHC, 1:1000 Western |
| Cardiac Troponin C (TNNC1) | 12G3, sc-52263, Santa Cruz | Mouse/IgG2b, monoclonal | 1:100 (1μg/mL) IHC, 1:100-1000 Western |
| Cardiac Troponin T (TNNT2) | 13-11, MA5-12960, Thermo Fisher | Mouse | 1:250 (2μg/mL) IHC, FAC, 1:500 Western |
| Myosin binding protein C (MYBPC3) | E-7, sc-137180, Santa Cruz | Mouse/IgG1, monoclonal | 1:100 (2μg/mL) IHC, 1:100-1000 Western |
| Human fibronectin HFN 7.1 | DSHB | Mouse, monoclonal | 1:250 (~2μg/mL) IF |
| WGA TRITC | W849, Invitrogen | | 1:100 (10 μg/mL) |
| WGA 350 | W11263, Invitrogen | | 1:100 (10 μg/mL) |
| Draq5 | 65-0880-92, Thermo Fisher | | 1:1000 (5μM IHC) |
| Phalloidin 488 | A12379, Thermo Fisher | | 1:100 (2unit/mL) |
| Hoescht | H21492, Thermo Fisher | | 1μg/mL |

**Supplemental Table 2: Primary antibodies and cellular counterstains used.**

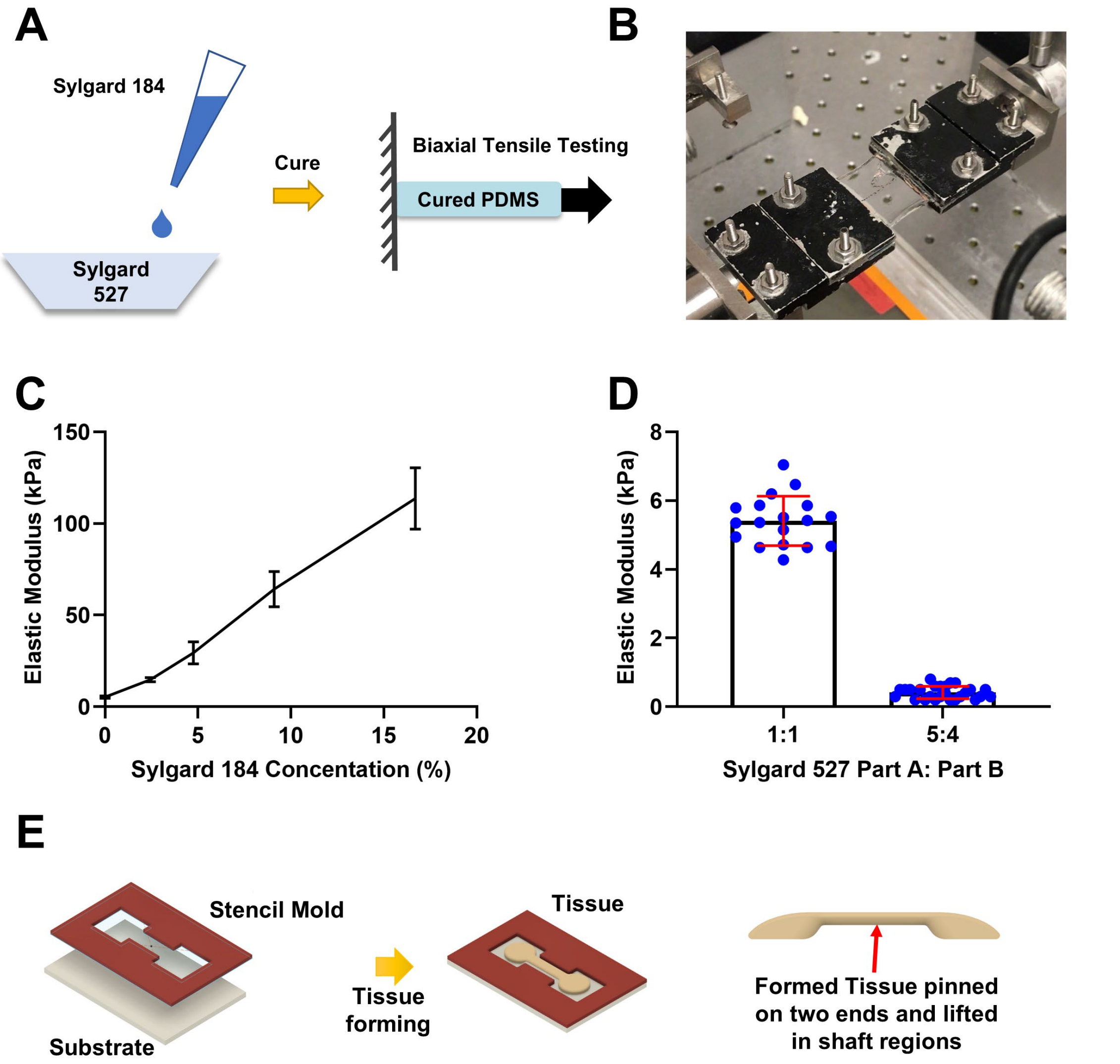


**Supplemental Fig. 1. Substrate characterization and tissue formation schematic. Controlling substrate mechanical properties using mix of sylgard 184 and sylgard 527. A) Blending different amounts of sylgard 184 to sylgard 527 to manipulate substrate stiffnesses. B) PDMS elastic modulus characterization through uniaxial tensile testing. C) Blended PDMS substrates can achieve tensile elasticity from 5 kPa up to 114 kPa. D) Ultra soft substrate can be achieved by blending different ratios of sylgard 527, part A to part B weight ratio of 5 to 4. E) Schematic of device fabrication and tissue formation[1].**

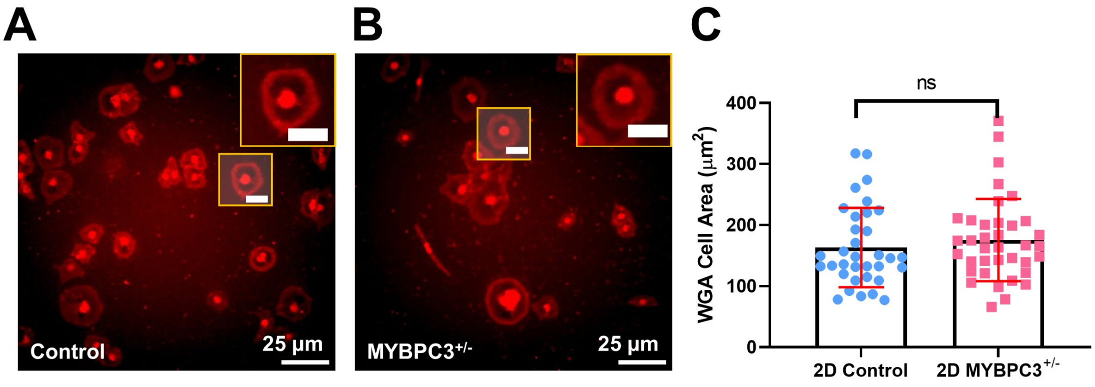


**Supplemental Fig. 2. Monolayer 2D iPSC-CM with MYBPC3$^{+/-}$ mutation does not exhibit cellular hypertrophy. (A) Representative WGA staining images of 2D control iPSC-CM. (B) Representative WGA images of 2D MYBPC3$^{+/-}$ iPSC-CM. (C) Quantitative analysis of cell area measured from both control and MYBPC3$^{+/-}$ cultures. *P* value 0.4336. n>5. Scale bar: 25 μm, insets 10 μm.**

| Tissue exclusion criteria based on tissue appearance | |
|---|---|
| “Good quality” tissue | 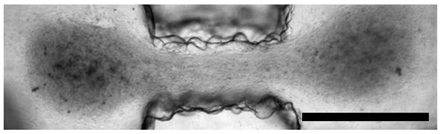 |
| Poor quality tissue type 1:<br>Visible cellular death and dark spots within the tissue | 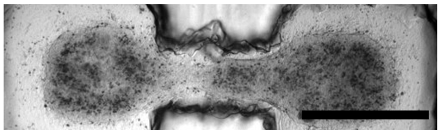 |
| Poor quality tissue type 2:<br>Partially detached tissue at the edges | 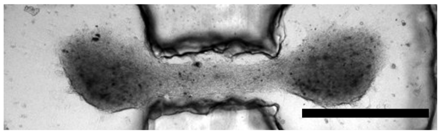 |
| Poor quality tissue type 3:<br>completely delaminated from substrate | 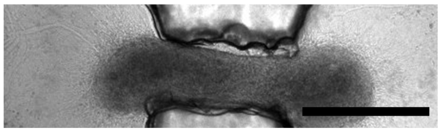 |

**Supplemental Fig. 3. Tissue exclusion criteria based upon tissue appearance. Within 75-95% cardiomyocytes batches, the lower quality tissues were excluded from physiological and electrophysiological studies. Only “good quality” tissues, which are uniformly distributed and compacted tissue with no obvious cellular death and detachment were selected for further analysis. We observe the inclusion/exclusion rate are not dependent on genotypes or stiffnesses.**

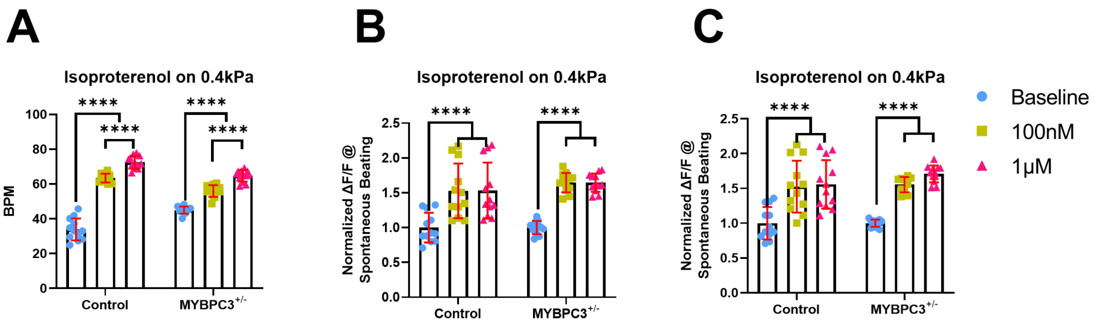


**Supplemental Fig. 4. Isoproterenol response for control and MYBPC3$^{+/-}$ tissues under soft 0.4 kPa condition. (A) Spontaneous beat rate changes for control and MYBPC3$^{+/-}$ 0.4 kPa tissues in response for isoproterenol. (b-C) Spontaneous (B) and 1.5 Hz (C) pacing $Ca^{2+}$ intake for control and MYBPC3+/- 15 kPa tissues in response for isoproterenol. **** indicates p value less than 0.05, 0.001 and 0.0001. Error bar: SD. n represents >3 individual differentiated tissue batches.**

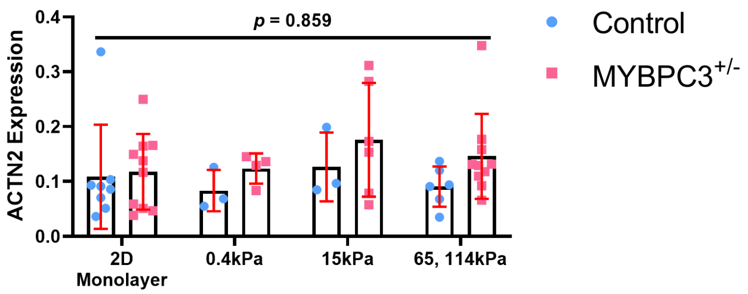


**Supplemental Fig. 5. Gene expression of ACTN2 expression between control and MYBPC3$^{+/-}$ at different environmental stiffnesses. Sarcomere actinin expression has no significant differences between conditions.**

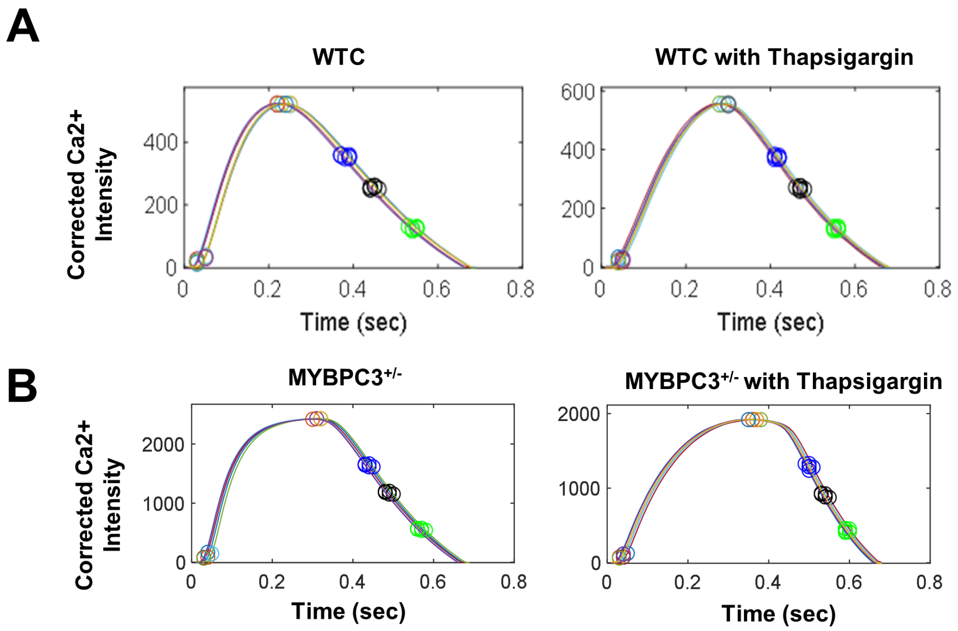


**Supplemental Fig. 6. SERCA inhibition using thapsigargin indicates SERCA inhibition causes the prolonged calcium upstroke for both genotypes, indicating both control and MYBPC3$^{+/-}$ tissues have functional SERCA. Meanwhile, SERCA inhibition does not recapitulate prolonged calcium plateau in control tissues.**

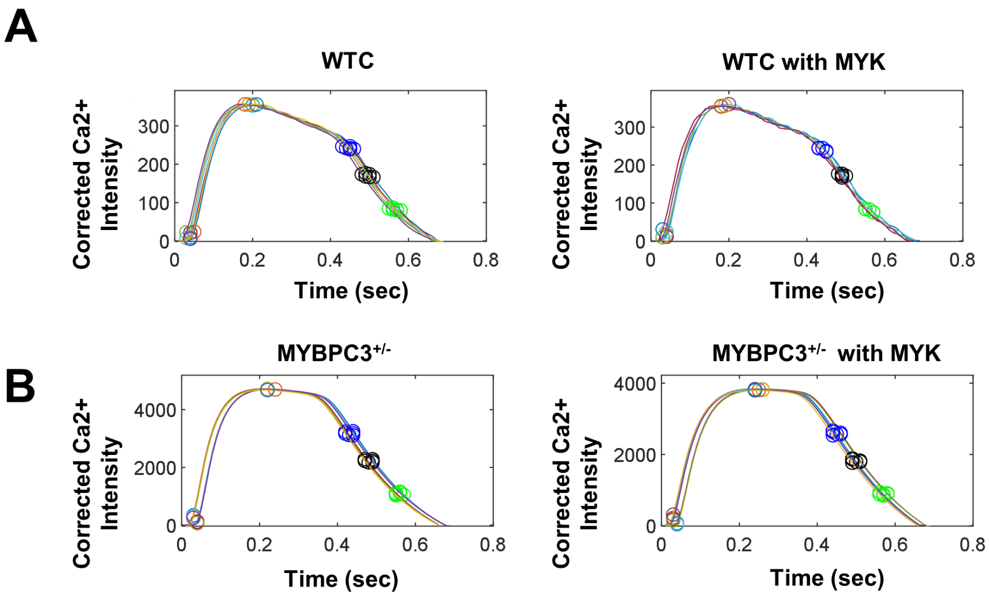


**Supplemental Fig. 7. Myosin inhibitor mavacamten (MYK) substantially reduced contractility of the μHM without effect calcium transient. A) Representative calcium transient of WTC 15 kPa tissue before and after treating with 0.5 μM MYK. B) Representative calcium transient of $MYBPC3^{+/-}$ 15 kPa tissue before and after treatment with 0.5 μM MYK.**

### References:

1. Guo, J. *et al.* Elastomer-Grafted iPSC-Derived Micro Heart Muscles to Investigate Effects of Mechanical Loading on Physiology. *ACS Biomater Sci Eng* (2020) doi:10.1021/acsbiomaterials.0c00318.
